# Competition and Synergy of Arp2/3 and Formins in Nucleating Actin Waves

**DOI:** 10.1101/2023.09.13.557508

**Authors:** Xiang Le Chua, Chee San Tong, X.J. Xŭ, Maohan Su, Shengping Xiao, Xudong Wu, Min Wu

## Abstract

The assembly and disassembly of actin filaments and their regulatory proteins are crucial for maintaining cell structure or changing physiological state. However, because of the tremendous global impact of actin on diverse cellular processes, dissecting the specific role of actin regulatory proteins remains challenging. In this study, we employ actin waves that propagate on the cortex of mast cell to investigate the interplay between formins and the Arp2/3 complex in the nucleating and turnover of cortical actin. Our findings reveal that the recruitment of FMNL1 and mDia3 precedes the Arp2/3 complex in cortical actin waves. Membrane and GTPase-interaction can drive oscillations of FMNL1 in an actin-dependent manner, but active Cdc42 waves or constitutively-active FMNL1 mutant can form without actin waves. In addition to the apparent coordinated assembly of formins and Arp2/3, we further reveal their antagonism, where inhibition of Arp2/3 complex by CK-666 led to a transient increase in the recruitment of formins and actin polymerization. Our analysis suggest that the antagonism could not be explained for the competition between FMNL1 and Arp2/3 for monomeric actin. Rather, it is regulated by a limited pool of their common upstream regulator, Cdc42, whose level is negatively regulated by Arp2/3. Collectively, our study highlights the multifaceted interactions, cooperative or competitive, between formins and Arp2/3 complex, in the intricate and dynamic control of actin cytoskeletal network.

## Introduction

Cells receive extracellular stimuli to trigger signal transduction pathways and activate actin cytoskeleton to regulate a plethora of diverse cellular processes, such as cell migration, adhesion, growth and division. Achieving precise regulation of these processes mandates the involvement of a repertoire of proteins capable of orchestrating actin assembly and turnover. Of particular significance are two major classes of actin nucleators: the Arp2/3 complex and formins (Campellone and Welch, 2010). While the simplistic view is that the Arp2/3 complex nucleates the side branching of actin filaments at 70° angle to allow the growth of new ‘daughter’ filaments, and formins are responsible for the elongation of the barbed ends of linear unbranched actin filaments, increasingly evident is the fact that these nucleators do not work in isolation within cells, but instead frequently engage in synergistic interactions, leading to the generation of higher order cellular architectures. While such interplay is well recognized in contexts such as leading-edge assembly during migration, immunological synapses, and adherens junctions (Chhabra and Higgs, 2007; Lappalainen et al., 2022), many mechanistic questions remain with regard to their assembly and crosstalks.

In recent years, the phenomenon of actin waves has gained widespread attention across diverse cell types, including but not limited to *Dictyostelium* (Bretschneider et al., 2009, 2004; Ecke et al., 2020; Gerisch et al., 2011; Jasnin et al., 2019; Schroth-Diez et al., 2009; Vicker, 2002), frog and echinoderm oocytes and embryos (Bement et al., 2015; Michaud et al., 2022; Swider et al., 2022; Tan et al., 2020; Wigbers et al., 2021), c elegans embryos (Michaux et al., 2018; Nishikawa et al., 2017; Yao et al., 2022), Drosohpilia embryo (Dehapiot et al., 2020), mouse embryos (Maître et al., 2015; Özgüç et al., 2022), mast cells (Su et al., 2020; Tong et al., 2023a, 2023b; Wu et al., 2013, 2018; Xiao et al., 2017; Xiong et al., 2016; Yang et al., 2017), Jurket T Cells (Lam Hui et al., 2014), neutrophils (Weiner et al., 2007), macrophage (Masters et al., 2016) and epithelial cells (Graessl et al., 2017; Mitsushima et al., 2010; Zhan et al., 2020). While supramolecular actin assemblies like lamellipodia and filopodia are dynamic, actin waves display cycles of actin assembly and disassembly in a much more coordinated and rapid fashion. A single cycle of assembly and disassembly takes place within a timeframe ranging from a few seconds to minutes (Wu et al., 2013), with the fastest oscillatory cycles with a mere 8 to10 seconds (Bretschneider et al., 2004; Ecke et al., 2020; Huang et al., 2013). The underlying rationale for inducing such transient yet recurrent patterns remains a subject of inquiry, with current literature implicating cortical actin waves in diverse cellular functions, including cell polarity and motility (Bolado-Carrancio et al., 2020; Fukushima et al., 2019; Li et al., 2020; Miao et al., 2019; Stankevicins et al., 2020), cell division (Bement et al., 2015; Dimitracopoulos et al., 2020; Flemming et al., 2020; Michaud et al., 2022; Xiao et al., 2017), cell growth (Huang et al., 2020), cell fate changes (Zhan et al., 2020), endocytosis (Yang et al., 2017), phagocytosis (Gerisch, 2010; Gerisch et al., 2009; Honda et al., 2021; Hörning et al., 2021; Lutton et al., 2023; Saito and Sawai, 2021) or exocytosis (Masters et al., 2016; Wollman and Meyer, 2012). In addition, intracellular actin waves have also been found on mitochondria, regulating membrane dynamics and partition during mitosis (Mitsushima et al., 2010; Moore et al., 2021, 2016). Despite certain similarities in the spatial and temporal patterns observed, it is pertinent to recognize the substantial differences characterizing actin waves across diverse systems. These waves are likely heterogeneous in nature and involve distinct higher order assembly of actin filaments, which remain largely uncharacterized.

In our current work, we employ the actin waves exhibited by mast cells as a model to investigate the nucleation mechanism for actin wave, in particular, how different actin nucleators collaborate within the same system. Our study unveils that both formins, FMNL1 and mDia3, are recruited to actin waves prior to the Arp2/3 complex with a consistent phase shift. The timing of formin recruitment to the membrane is set by its interaction with GTPase, not with actin. Strikingly, constitutively-active mutant of FMNL1 can propagate waves even in an F-actin-depleted state. Lastly, we found that acute inhibition of Arp2/3 complex enhance formin recruitment. This effect does not arise due to the competition for the actin monomer, rather, it is due to the availability of upstream regulators, which is negatively regulated by Arp2/3. Collectively, our analysis reveals the intricate interplay between Arp2/3 and formin-associated actin polymerization in modulating actin collective dynamics through mechanisms involving both cooperation and competition.

## Results

### Arp2/3 complex and formins FMNL1 and mDia3 assemble as coordinated waves

Our previous studies revealed the propagation of cortical oscillatory waves in resting and stimulated rat basophilic leukemia (RBL-2H3) cells, as well as the recruitment of FMNL1, mDia3 and FHDC1 to the ventral actin waves that are Cdc42-dependent (Tong et al., 2023a). These three actin nucleators belong to the family of 15 formins, a major class of actin polymerizing proteins responsible for the elongation of the fast-growing, barbed ends of linear unbranched actin filaments. How they compare to Arp2/3, which generated branched network, is not clear. Here we examined the dynamics of these actin nucleators, first individually then in combination. When imaged by total internal reflection microscopy (TIRFM), GFP-Arp3 appeared as distinct clusters of puncta that exhibited sequential assembly and disassembly cycles (n= 21 cells) (**Figure 1A**, **Video 1**). These recurrent cycles of assembly and disassembly were phase-shifted at different location and resembled wave-like movement but the individual loci did not move in its lifetime. With the maximum intensity projections constructed from the timelapse, we found out that the probablity of Arp2/3 puncta assembles were relatively uniform throughout the cell, except that they were signficantly reduced at the peripheral edges of the cell (**Figure S1A**). The assembly of Arp2/3 (GFP-Arp3) preceded F-actin (LifeAct-mRuby) by approximately 2 sec in cortical waves (n= 28 cells, 10 experiments) (**Video 1**). To ensure an unbiased analysis of these waves, we generated kymographs at three randomly selected regions within a cell expressing GFP-Arp3 (**Figure S1B**). At these chosen regions, we also plotted the changes in fluorescence intensity using three different ROI sizes (5 pixel x 5 pixel, 20 x 20 pixel and 40 x 40 pixel) (**Figure S1C**). Our analysis showed that the observed wave cycles in the fluorescence intensity plots remained consistent across the three different ROI sizes, as judged by the similar patterns of the rise and decay of each peak. However, peaks of intensity changes acquired from larger ROI size appeared to be smoother than smaller ROI sizes, due to the averaging effect. The number of peaks observed in the plots corresponds to the number of bands in the kymograph re-slices. To maintain consistency, we used an ROI size of 20 pixel x 20 pixel (equivalent to 2.2 µm x 2.2 µm) for all our data analysis in this study.

**Figure 1.**
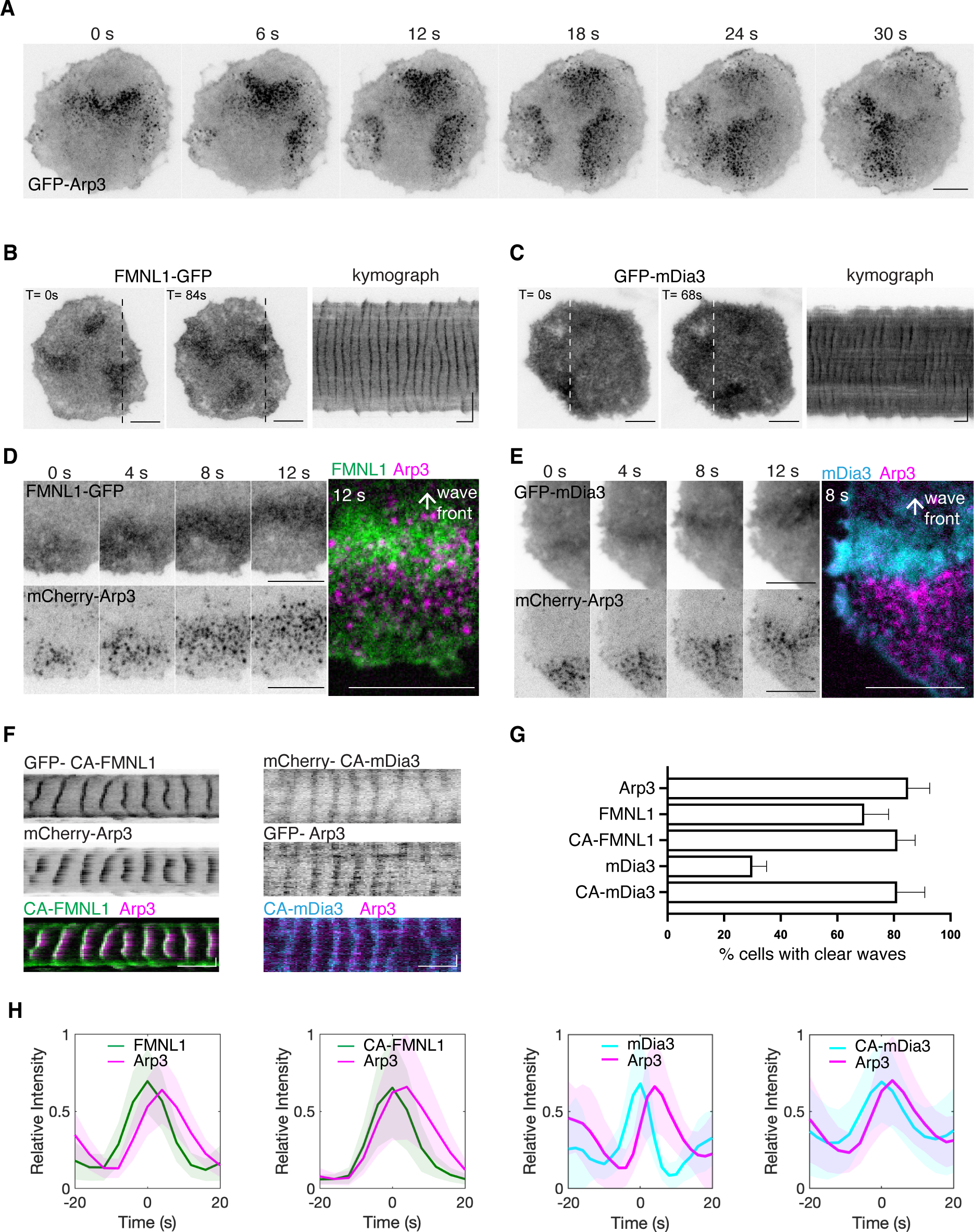
Localization of Arp3, FMNL1 and mDia3 in cortical traveling waves. **(A)** Representative TIRFM time-lapse montage of six frames (6s interval) of a cell expressing GFP-Arp3. **(B-C)** Representative micrographs and kymographs of a cell expressing **(B)** FMNL1-GFP and **(C)** GFP-mDia3. **(D-E)** Sequential montage of four frames (4s interval) of a region of a representative cell co-expressing **(D)** FMNL1-GFP and mCherry-Arp3, **(E)** GFP-mDia3 and mCherry-Arp3, followed by a two-color merge micrograph showing the wave front of **(D)** FMNL1-GFP (green) and mCherry-Arp3 (magenta) and **(E)** GFP-mDia3 (cyan) and mCherry-Arp3 (magenta). **(F)** Representative kymographs of a cell co-expressing GFP-CA-FMNL1 (green) and mCherry-Arp3 (magenta), mCherry-CA-mDia3 (cyan) and GFP-Arp3. **(G)** Quantification for the percentage of cells exhibiting clear waves of different formin constructs from unbiased selection exp (> 11 cells from 2-3 independent experiments; n= 85 % of cells exhibiting clear waves of either GFP-Arp3 or mCherry-Arp3 in 3 exp; n= 69.5 % of cells exhibiting clear waves of FMNL1-GFP or FMNL1-mCherry in 3 exp; n= 81.3 % of cells exhibiting clear waves of GFP-CA-FMNL1 in 2 exp; n= 30 % of cells exhibiting clear waves of GFP-mDia3 or mCherry-mDia3 in 3 exp; n= 81.2 % cells exhibiting clear waves of mCherry-CA-mDia3 in 2 exp). **(H)** Fluorescence intensity profiles of GFP- or mCherry-Arp3 (magenta) aligned with respect to FMNL1-GFP (green), GFP-CA-FMNL1 (green), GFP-mDia3 (cyan), mCherry-CA-mDia3. All greyscale micrographs and kymographs are shown in inverted lookup table. Horizontal scale bars in micrographs: 10 µm. Vertical scale bars in kymographs: 10 µm. Horizontal scale bars in kymographs: 1 min.

Unlike the distinct Arp3 clusters, FMNL1 waves appeared as diffusive but visually apparent cloud (n= 33 cells, 13 experiments) (**Figure 1B**). Compared to FMNL1, mDia3 waves had only subtle increase in intensity over the background levels, resulting in a less distinct contrast (n= 28 cells, 10 experiments) (**Figure 1C**). To confirm that mDia3 waves were a result of its recruitment to the plasma membrane, not simply an artefact caused by membrane fluctuations that could lead to changes of TIRF signals, we co-expressed GFP-mDia3 with a plasma membrane marker (PM-mScarlet). In cells that exhibited waves of mDia3, we did not observe intensity oscillations of plasma membrane marker at the same amplitude and regularity, indicating that the observed waves of formins could not be caused by membrane lifting from the TIRF plane (n= 12/12 cells in 2 experiments) (**Figure S2**).

To gain an understanding of the nucleation process driving the actin wave formation, we visualized the dynamics between these two classes of nucleators by expressing them in the same cell. We co-expressed the fluorescently-labelled marker for FMNL1 or mDia3, with mCherry-Arp3. We found that waves of FMNL1 and Arp3 always appear in the same cells and were coordinated (**Figure 1D**, **Video 2**). In these cells, waves of Arp3 trailed behind the corresponding waves of FMNL1 by about 2 sec (n= 29 cells, 11 experiments) (**Figure 1D**). In contrast, mDia3 only appeared as coordinated waves with Arp3 in a subset of the cells (**Figure 1E**). Given the auto-regulatory properties of diaphanous-related formins, we also tested the effect of releasing formins from their autoinhibited states. The L1062D mutation at the DAD motif of FMNL1 prevents its interaction with the DID motif, thus rendering it constitutively activate (Seth et al., 2006). For mDia3, we deleted the DAD domain to generate a constitutively-active (CA) mutant of mDia3. We co-expressed CA mutants of mDia3 (mCherry-CA-mDia3) and Arp3 (GFP-Arp3) in cells and observed increased appearance of coordinated waves of CA-mDia3 and Arp3 (n= 11 cells, 2 experiments) (**Figure 1F**). CA-FMNL1 also displayed a modest increase of the percentage of cells showing regular oscillation (n= 9 cells, 3 experiments). We quantified fluorescent intensity using Fast Fourier Transform (FFT) and used the presence of dominant frequency peak as a criteria, we determined that wild-type FMNL1, CA-FMNL1, wild-type mDia3 and CA-mDia3 displayed oscillatory wave patterns in 69.5 %, 81.3 %, 30 % and 81.2 % of the cells counted respectively (**Figure 1G**). Intensity profiles acquired from co-expression of Arp3 with either wild-type or CA formins showed that CA mutants shared similar phase as wild-type proteins, and that FMNL1, CA-FMNL1, mDia3 and CA-mDia3 were all recruited ahead of Arp3 in cortical wave propagation (FMNL1 precedes Arp3 by ∼1.9 sec; CA-FMNL1 precedes Arp3 by ∼1.8 sec; mDia3 precedes Arp3 by ∼3.8 sec; CA-mDia3 precedes Arp3 by ∼2.1 sec) (**Figure 1H**). Our results collectively illustrate the variations in the distributions of actin nucleators. Nonetheless, the recruitment of formins (FMNL1 and mDia3) preceding Arp2/3 complex was robustly seen.

### FMNL1 recruitment to waves requires the presence of a functional GBD and the alleviation of auto-inhibition interactions

To gain further insights of how FMNL1 was recruited to the membrane, we co-expressed a range of truncated and point mutants of FMNL1 (Seth et al., 2006), together with Arp3. FMNL1 share the conserved domain architecture with the diaphanous family of formins, defined by their N-terminus GTPase binding domain (GBD), followed by the diaphanous inhibitory domain (DID), coiled-coil (CC), dimerization domain (DD), formin homology-1 (FH1) and 2 (FH2) domains, and finally, the diaphanous auto-regulatory domain (DAD) at the C-terminus. While FMNL1-GFP and GFP-CA-FMNL1 were recruited before Arp3 and did not co-cluster with Arp3 (**Figure 2A**, **Video 2**, **Video 3**), the C-termini mutant of FMNL1 with the FH2 and DAD motifs (GFP-FMNL1CT) appeared as punctate structures (**Figure 2B**, **Video 4**). These punctate of FMNL1CT co-localized spatially and temporally with the Arp3 puncta (n= 12/12 cells, 2 experiments). Waves exhibited by the FMNL1CT were highly irregular and propagated only in localized regions of the cell, in contrast with cells co-expressing mCherry-Arp3 with GFP-CA-FMNL1 (**Figure 2C**, **2D**). These results suggest that C-termini of FMNL1 and how it interacts with actin unlikely explain the early recruitment of FMNL1.

**Figure 2.**
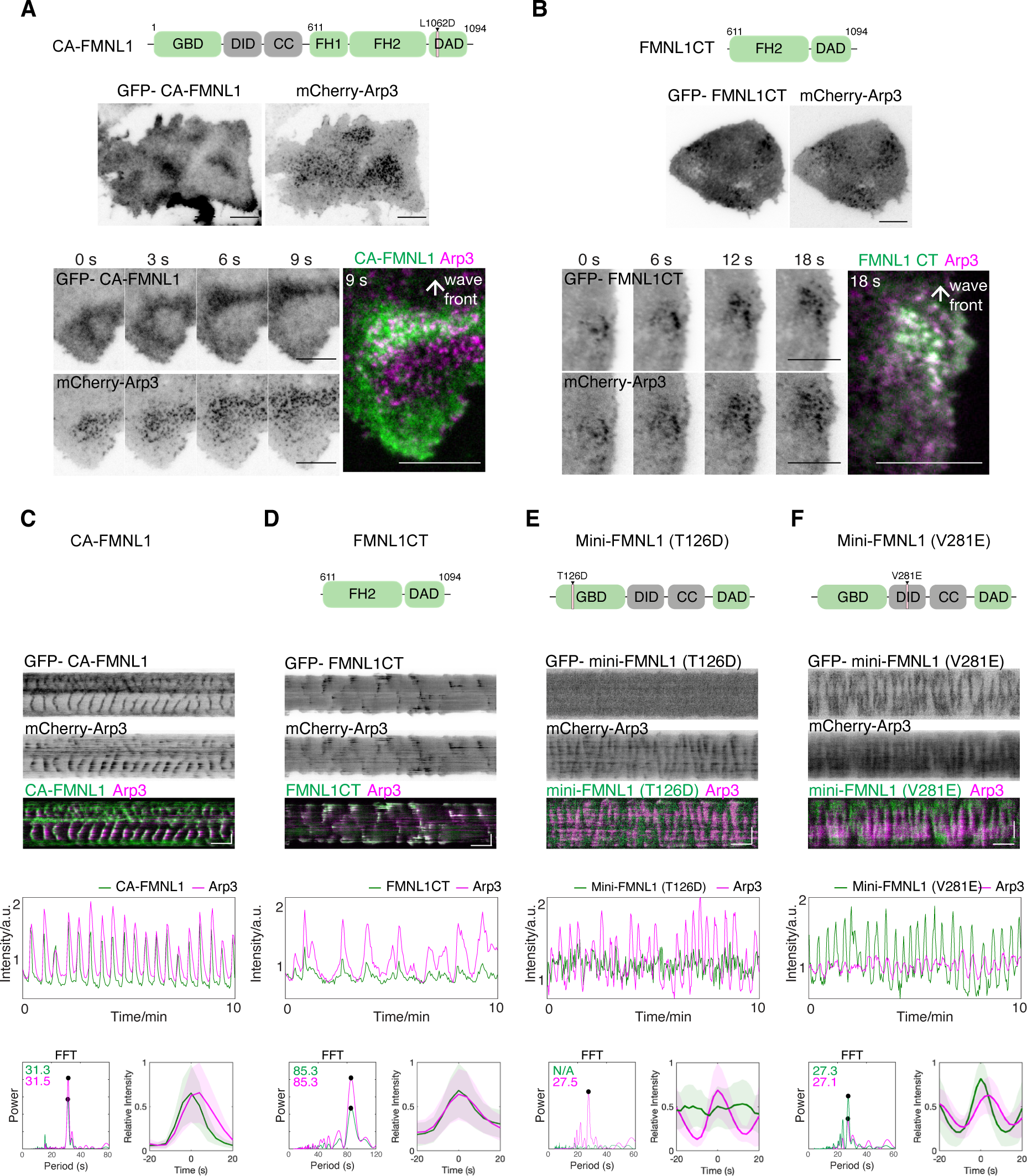
GTPase-binding domain and relieved auto-inhibition are necessary for FMNL1 assembly in waves. **(A-B)** Domain schematics of CA- and C-termini-mutants of FMNL1 used in our study. Representative micrographs and sequential montage of four frames (3-6s interval) of a region of a cell co-expressing mCherry-Arp3 (magenta) and **(A)** GFP-CA-FMNL1 (green) or **(B)** GFP-FMNL1CT (green). **(C)** Representative kymographs, intensity plot and FFT profile of a cell co-expressing GFP-CA-FMNL1 (green) and mCherry-Arp3 (magenta). **(D-F)** Domain schematics of truncated and point mutants of FMNL1 used in our study. Representative kymographs, intensity and FFT profiles of a cell co-expressing mCherry-Arp3 (magenta) and the **(D)** C-termini-, **(E)** mini-FMNL1 truncated of FH1-FH2 with T126D mutation at GBD motif-, or **(F)** mini-FMNL1 truncated of FH1-FH2 with V281E mutation at DID motif-mutant of FMNL1. FMNL1 mutants are false-colored green. Representative intensity profiles of mCherry-Arp3 (magenta) aligned with respect to FMNL1 mutants (green). All greyscale micrographs and kymographs are shown in inverted lookup table.

We next conducted experiments using two “mini” mutants of FMNL1 whose FH1-FH2 domains were substituted by a (Gly-Gly-Ser)_2_ linker (Seth et al., 2006). The first mutant, mini-FMNL1-T126D, is deficient in its binding to its endogenous activating GTPase, and the second mutant, mini-FMNL1-V281E, is deficient in DAD binding and mimicks an active conformation (Seth et al., 2006). GFP-mini-FMNL1-T126D did not exhibit any observable wave-like behavior, despite of the clear wave of Arp3 co-expressed in the same cell (n= 0/6 cells) (**Figure 2E**). GFP-mini-FMNL1-V281E was observed to propagate clear waves (n= 7/7 cells) (**Figure 2F**). Waves of mini-FMNL1-V281E mutant precede the Arp3 waves (**Figure 2F**), similar to the temporal dynamics exhibited by wildtype FMNL1 or CA-FMNL1. However, the amplitude of Arp3 waves were reduced (**Figure 2F**). Taken together, these results illustrate the order in which the membrane and actin interaction sites orchestrate the temporal recruitment and action of FMNL1. GTPase-interaction domain is not only essential for FMNL1’s recruitment to the membrane, but is also responsible for its timing. In the absence of the GTPase-binding domain, the expression of the FH2 and DAD motifs of FMNL1 alone allows its recruitment to wave, but it results in altered phase, localization and wave geometry.

### Membrane waves of active Cdc42 and constitutively-active FMNL1 continue to exist in cells even in the absence of F-actin

If membrane binding determines the phase of FMNL1 recruitment, we next wonder whether FMNL1 could oscillate without actin oscillations. We sought to create experimental conditions that would reduce or eliminate actin oscillation. To achieve this, we used cytochalasin-D, a pharmacological inhibitor that disrupts the actin cytoskeleton by blocking the barbed ends of actin filaments (Hetrick et al., 2013). We used cells expressing a Cdc42 biosensor (Cdc42 BD-GFP), which served as an indicator for membrane waves, LifeAct-mRuby, which serves as as a control marker for actin. Through systematic experimentation of varying drug doses and incubation time, we found that chronic treatment with cytochalasin-D (2-4 µM) over a period of 18-24 hours could uncouple membrane oscillations from actin oscillations (**Figure 3A**). We could robustly observe cells exhibiting clear waves of active Cdc42 in the absence of detectable LifeAct waves (n= 60/60 cells in 7 experiments) (**Figure 3B**, **Video 5**). Actin waves gradually resumed if cytochalasin treated cells were allowed to recover in fresh growth media. The observed waves of Cdc42 were robust but oscillatory periodicities were longer compared to waves in untreated conditions (**Video 6**). Under this condition, GFP-CA-FMNL1 exhibited clear waves in the absence of LifeAct-mRuby waves (n= 3/3 cells, 2 experiments) (**Figure 3C**, **Video 7**). On the other hand, waves of Arp3, FMNL1 or mDia3 were all abolished, when LifeAct waves were absent, and all could resume in the recovery phase together (n= 20 cells for mCherry-Arp3 and mNG-LifeAct in 2 experiments; n= 7 cells for FMNL1-GFP and LifeAct-mRuby in 2 experiments; n= 11 cells for GFP-mDia3 and LifeAct-mRuby in 2 experiments) (**Figure 3D-F**). Collectively, our results showed that the propagation of Cdc42 and CA-FMNL1 waves can be independent of actin waves. While membrane-dependent interactions are sufficient to drive CA-FMNL1 recrutiment and oscillations, this interaction is not strong enough for wildtype proteins which likely exist partially in its auto-inhibited state.

**Figure 3.**
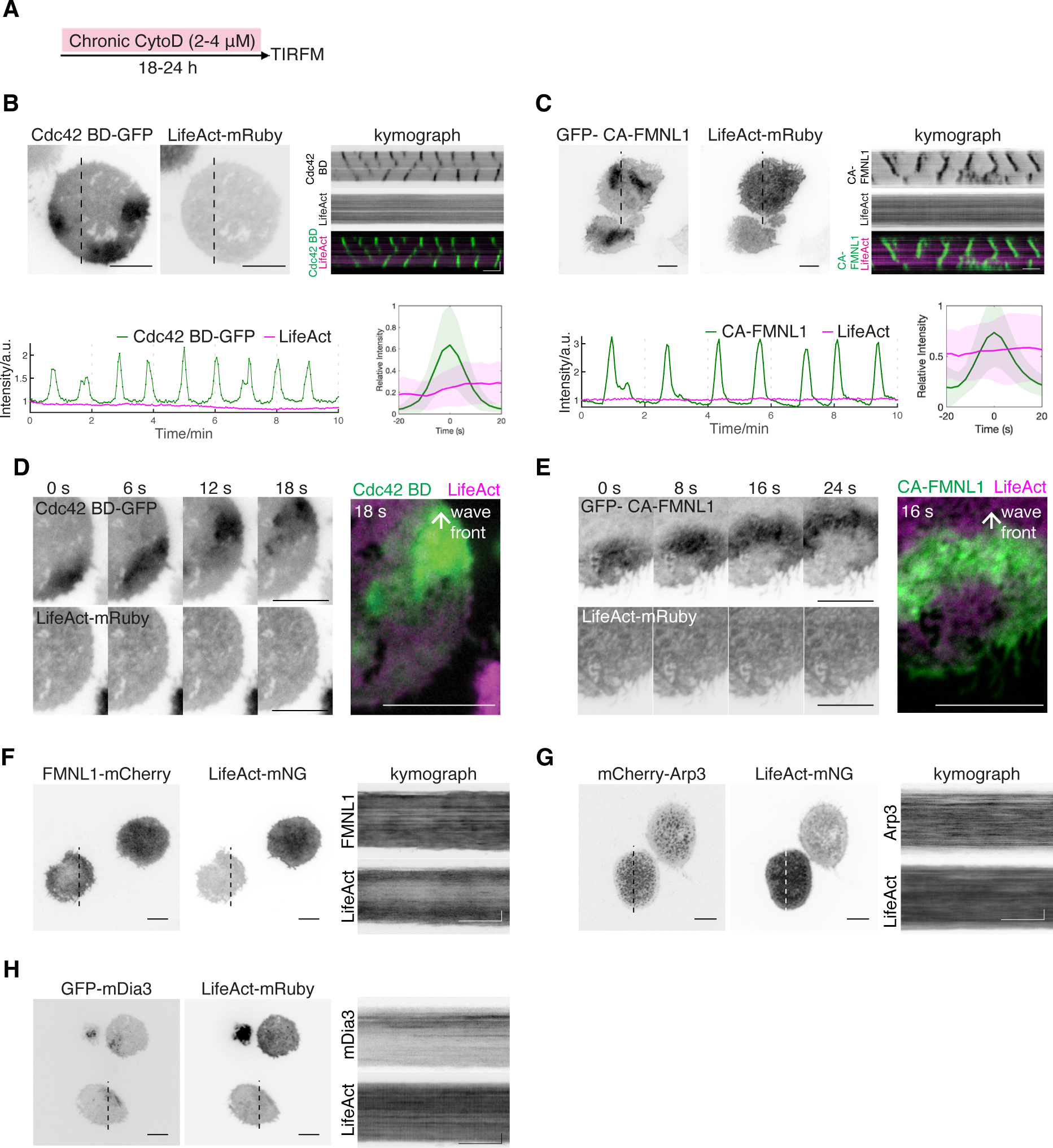
Cortical waves of active Cdc42 and constitutively-active FMNL1 persist in actin-depleted state. **(A)** Treatment schematic for the chronic depletion of F-actin with cytochalasin-D. **(B)** Representative micrographs and kymographs of a chronically pre-treated cell stably-expressing Cdc42 BD-GFP (green) co-transfected with LifeAct-mRuby (magenta). Representative intensity plot and profile of Cdc42 BD-GFP (green) and LifeAct-mRuby (magenta). **(C)** Representative greyscale micrographs and kymographs of a chronically-treated cell co-expressing GFP-CA-FMNL1 (green) and LifeAct-mRuby (magenta). Representative intensity plot and profile of GFP-CA-FMNL1 (green) and LifeAct-mRuby (magenta). **(D-F)** Representative micrographs and kymographs of chronically pre-treated cells co-expressing LifeAct-mRuby or LifeAct-mNG and **(D)** FMNL1-mCherry, **(E)** mCherry-Arp3, or **(F)** GFP-mDia3 (n= 0/3 cells exhibiting waves of FMNL1-GFP and LifeAct-mRuby in 2 exp; n= 0/18 cells exhibiting waves of mCherry-Arp3 and LifeAct-mNG in 2 exp; n= 0/13 cells exhibiting waves of GFP-mDia3 and LifeAct-mRuby in 2 exp). All greyscale micrographs and kymographs are shown in inverted lookup table. Horizontal scale bars in micrographs: 10 µm. Vertical scale bars in kymographs: 10 µm. Horizontal scale bars in kymographs: 1 min.

### Competitive assembly between the Arp2/3 complex and FMNL1/mDia3

To investigate the potential role of Arp2/3 that follows formins in polymerizing actin waves, we used CK-666, a pharmacological inhibitor of the Arp2/3 complex (Hetrick et al., 2013). When CK-666 was applied at concentrations ranging from 50-150 µM, most of the mCherry-Arp3 puncta disappeared along with the Arp3 oscillations (**Figure 4A**). Surprisingly, the intensities of FMNL1-GFP waves were enhanced (**Figure 4A**). Moreover, we also observed an increase in the oscillatory periodicities, as indicated on the wavelet analysis (50 CK-666: n= 9 cells, 150 µM CK-666: n=3 cells) (**Figure 4B**, **Video 8**). Similar antogonistic relationship was found between GFP-mDia3 and mCherry-Arp3 (50 µM CK-666: n= 7 cells, 150 µM CK-666: n= 7 cells) (**Figure 4C-D**, **Video 9**). Under these conditions when Arp3 puncta and waves were perturbed after treatment with CK-666 at 50 µM, we saw an increase in the intensities of F-actin waves (LifeAct-mRuby) (n= 9/9 cells, 4 experiments) (**Figure 4E-F**). At a higher dosage of CK-666 at 200 µM, we observed inhibition of both LifeAct and Arp3 waves (n= 2 cells, 2 experiments) (**Figure 4F**). Further increase in the concentration of CK-666 to 250 µM completely abolished both LifeAct and Arp3 waves (n= 3 cells, 2 experiments) (**Figure 4G**). Taken together, our results reveal an increase in formin recruitment and actin polymerization in response to intermediate doses of Arp2/3 inhibition by CK-666. Arp2/3 complex, which was recruited after formins, appear to limit formin-mediated actin polymerization in actin waves.

**Figure 4.**
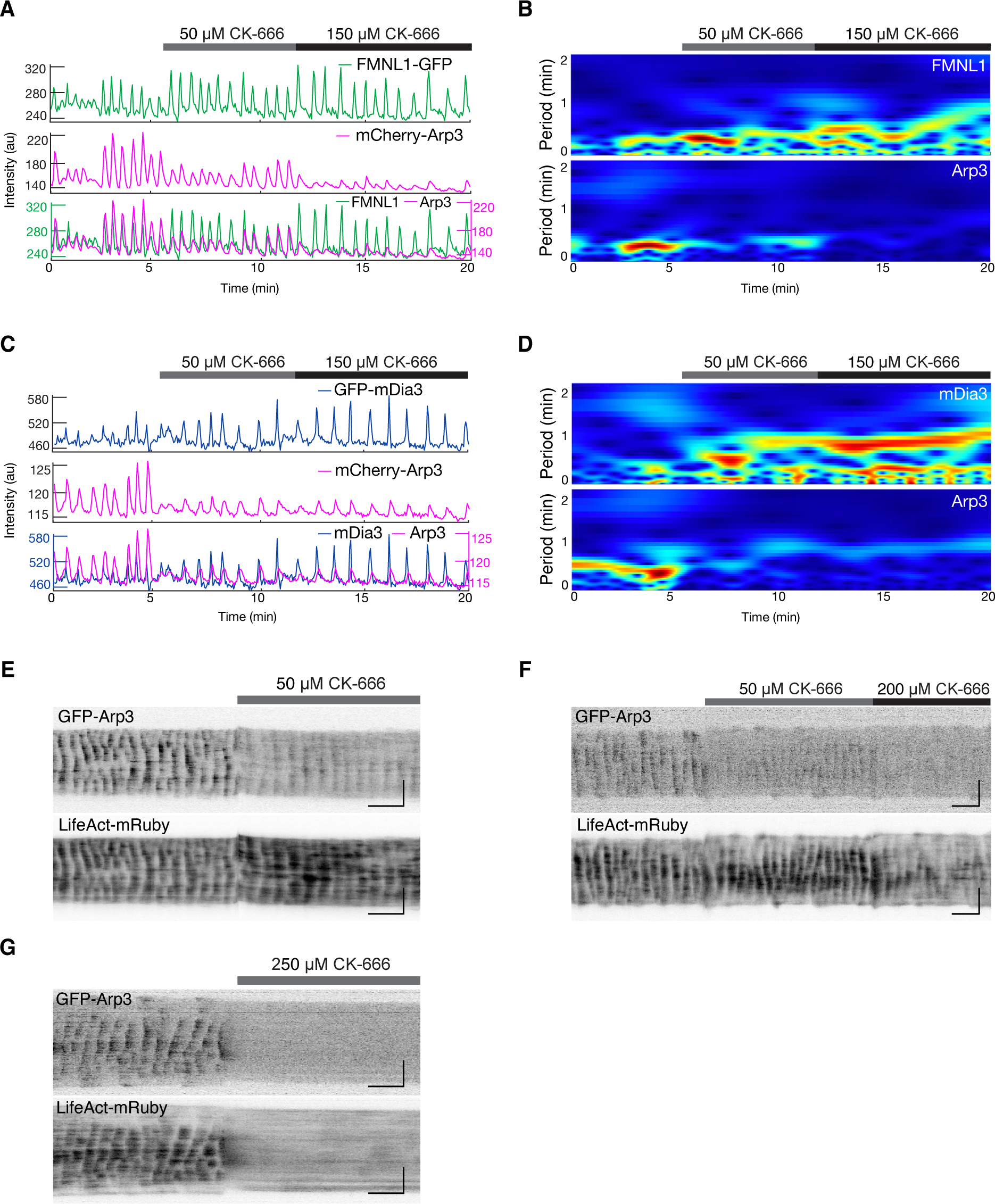
Enhanced formin-mediated actin polymerization in response to Arp2/3 inhibition. **(A-B)** Representative **(A)** intensity plots and **(B)** wavelet analyses of a cell co-expressing FMNL1-GFP (green) and mCherry-Arp3 (magenta) before and after treated with 50 µM, followed by 150 µM CK-666. **(C-D)** Representative **(C)** intensity plots and **(D)** wavelet analyses of a cell co-expressing GFP-mDia3 (blue) and mCherry-Arp3 (magenta) before and after treated with 50 µM, followed by 150 µM CK-666. **(E)** Representative kymographs of a cell co-expressing GFP-Arp3 and mCherry-Arp3 before and getting treated by 50 µM CK-666. **(F)** Representative kymographs of a cell co-expressing GFP-Arp3 and mCherry-Arp3 before and getting treated by 50 µM, followed by 200 µM CK-666. **(G)** Representative kymographs of a cell co-expressing GFP-Arp3 and mCherry-Arp3 before and getting treated by 250 µM CK-666. Horizontal scale bars in micrographs: 10 µm. Vertical scale bars in kymographs: 10 µm. Horizontal scale bars in kymographs: 1 min.

### FMNL1 and Arp2/3 competition through a limited pool of active Cdc42

The enhanced formin recruitment upon Arp2/3 inhibition implies that the recruitment of formins under normal conditions is competitively inhibited by Arp2/3. The prevailing model proposes a competition between the Arp2/3 complex and formins due to a limiting pool of actin monomer (Burke et al., 2014; Henty-Ridilla and Goode, 2015; Rotty et al., 2015; Rotty and Bear, 2014; Suarez et al., 2015; Vitriol et al., 2015). However, our data do not align with this model in its simplest form. First, limited pool of actin monomer can only explain a competition between Arp2/3 and formin activity, but not their localization and amplitude of recruitment. Both formin were recruited before Arp3 and F-actin with a significant time shift. Specifically, FMNL1 precedes Arp3 by approximately 1.9 sec (n= 29/29 cells, 11 experiments), while FMNL1 precedes LifeAct by approximately 4.5 sec (n= 5/5 cells, 3 experiments). The rate of FMNL1 recruitment also did not slow down when Arp was recruited (**Figure 1H**). Second, when oscillations of FMNL1-GFP and mCherry-Arp3 were monitored over time (ranging between 30-60 minutes), we consistently observed coordinated amplitudes of FMNL1 and Arp3 oscillations (n= 14 cells, 8 experiments) (**Figure 5A**), suggesting that it is possible to increase both Arp3 or FMNL1 recruitments from one cycle of 20 sec to the next 20 sec. If recruitment of one nucleator is competiting with another one, one would expect to see opposite amplitude for formin and Arp2/3 oscillations. These similar trends in the oscillations of both nucleators over many cycles indicate that the limiting factor dictating the competition, as reflected in the acute Arp2/3 inhibitor experiment, must be short-lived and does not carry over to the next cycle.

**Figure 5.**
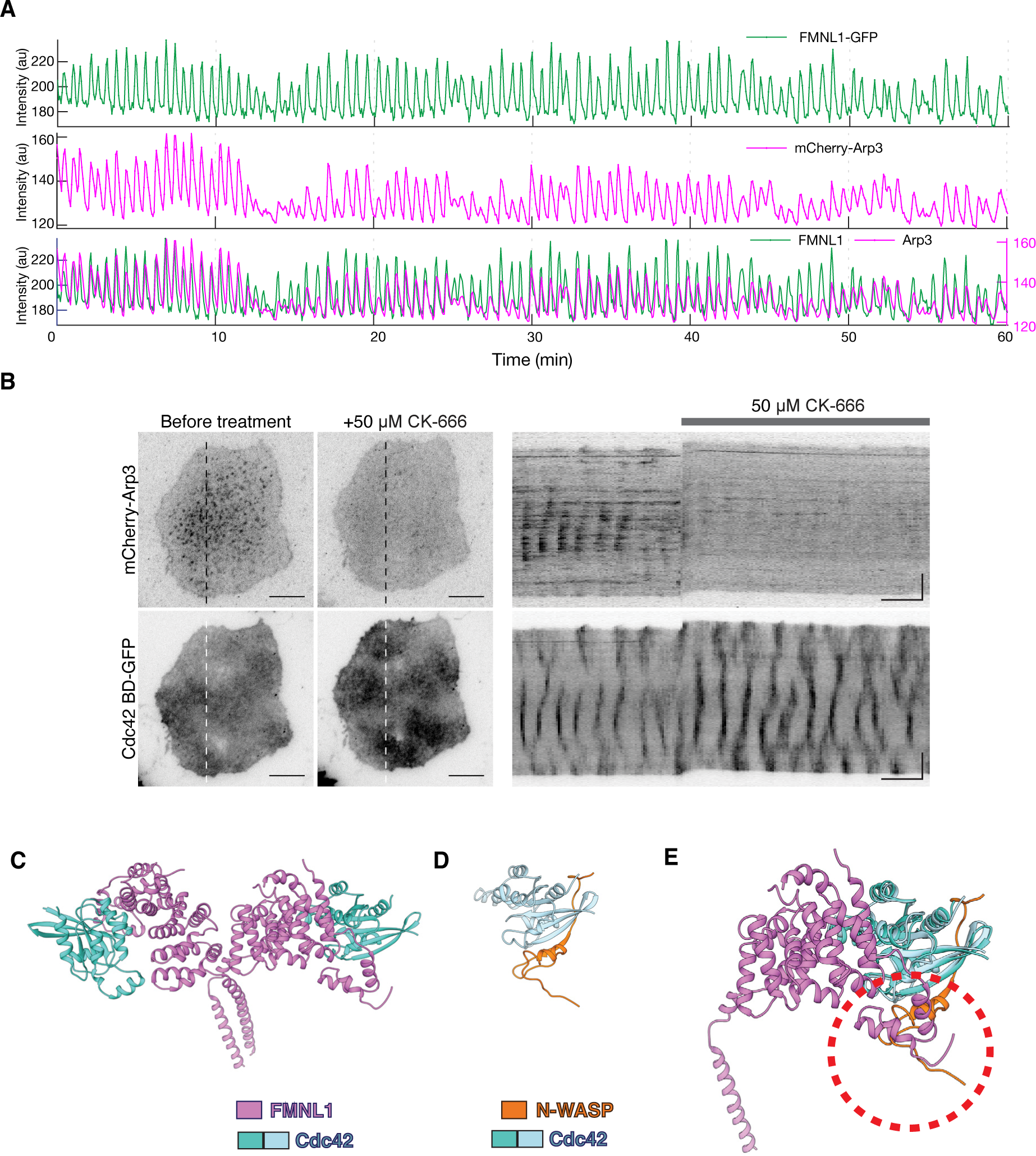
FMNL1 and Arp2/3 complex compete for upstream Cdc42 GTPase. **(A)** Representative intensity profile of a cell co-expressing FMNL1-GFP (green) and mCherry-Arp3 (magenta) over an acquisition period of 1 h (n= 14 cells co-expressing clear waves of FMNL1-GFP and mCherry-Arp3 acquired between 20 – 60 min in 8 exp). **(B)** Representative micrographs and kymographs of a cell stably-expressing Cdc42 BD-GFP co-transfected with mCherry-Arp3 before, and after treatment with 50 µM CK-666. **(C)** Left: Structure of the dimeric Cdc42/FMNL1 complex (pdb: 4ydh). **(D)** Structure of the Cdc42/N-WASP complex (pdb: 1cee) **(E)** Structure of the Cdc42/N-WASP complex aligned with the structure of monomeric Cdc42/FMNL1 complex on the basis of Cdc2 using Chimera. Models of Cdc42 are shown in light blue and turquoise, model of the GTPase binding domain of N-WASP is shown in orange, and the model of the N-terminal domain of FMNL1 is shown in purple. Red dashed circle shows the region where FMNL1 and N-WASP will clash.

We therefore speculate that the competitive assembly between FMNL1 and the Arp2/3 complex is due to a limited pool of upstream regulator, which is dynamically set within a given cycle. We examined the behavior of Cdc42, a shared upstream regulator for FMNL1 and Arp2/3. Cdc42 GTPase regulates Arp2/3-mediated actin polymerization through the activation of N-WASP/WASP, which subsequently activates the Arp2/3 complex. Furthermore, previous studies have shown that Cdc42 can form complexes with both WASP and FMNL1 (Abdul-Manan et al., 1999). By treating cells co-expressing Cdc42 BD-GFP and mCherry-Arp3 with 50 µM CK-666, we found out that Cdc42 activity was enhanced (n= 6/6 cells, 2 experiments) (**Figure 5B**, **Video 10**). These results suggest an apparent negative feedback from Arp2/3 to Cdc42. Furthermore, an analysis of the available structures of the Cdc42/WASP and Cdc42/FMNL1 complexes revealed significant steric clashes between these two complexes (**Figure 5C-E**). Taken together, our quantitative data are consistent with a model where the antagonism between Arp2/3 complex and formins is regulated more upstream, likely involving their competition for Cdc42.

## Discussion

In this paper, we characterized in detail the interplay between Arp2/3 and formins in actin waves in mast cells. These collective dynamcis of actin waves are ideal experimental systems. Detection of formin activity in living cells often require the utilization of constitutive active mutants, which may perturb the physiological cytoskeletal network and actin dynamics. Direct visualization of these two groups of nucleators in the same physiological condition has been challenging. Genetic perturbations would be too slow and were limited in dissecting systems with multiple levels of positive and negative feedbacks. Here, we were able to visualize the dynamics of these important actin regulators at physiological conditions with high spatial and temporal resolution, which allowed us to quantitaively characterize their interplay, including both coordination and competition.

### Role of Arp2/3 in actin waves

Our data do not argue against an essential role of Arp2/3 in nucleating actin waves. The recruitment of Arp2/3 to Cdc42 waves were the most robust. They appear in every single cell with Cdc42 and actin waves. In contrast, FMNL1 waves appear in the majority of cells with Cdc42 waves, while mDia3 appear in a subset of cells with Cdc42 waves. Nevertheless, the relative minor role Arp2/3 plays in Cdc42-dependent actin waves is surprising, since Arp2/3 is the most well-established downstream effector for Cdc42 and N-WASP. Activating actin polymerization by Cdc42 through N-WASP and Arp2/3 are classic pathways that have been extensively investigated (Machesky and Insall, 1998; Miki et al., 1998). There studies demonstrated both necessity and sufficiency of Arp2/3 in Cdc42-regulated actin polymerization. Necessity was proved by depleting Arp2/3 (Mullins and Pollard, 1999) or N-WASP in the extract-based reconstitution assays for Cdc42-dependent actin polymerization (Rohatgi et al., 2000, 1999), and sufficiency was shown by in vitro actin polymerization assays using purified Arp2/3 with N-WASP (Rohatgi et al., 1999) or SCAR (Machesky et al., 1999). These interaction of Cdc42 and WASP are also supported by structural evidence (Abdul-Manan et al., 1999; Kim et al., 2000). On the other hand, Arp2/3 was not identified through an unbiased screen for actin nucleator. Data derived from Arp2/3 knockout cells have suggested that the Arp2/3 complex is not as ubiquitously essential for actin dynamics as originally thought (Rotty et al., 2017; Suraneni et al., 2012; Wu et al., 2012).

Our data showing the relative minor contribution of Arp2/3 in the actin waves support the emerging view that Arp2/3 nucleated actin branches could be more specialized. Within the single cell, Arp2/3 is preferentially localized in the cell center relative to the cell edge. Actin waves after Arp2/3 inhibition persisted, but have longer cycle time. These suggest that the function of Arp2/3 could be to promote faster turnover of actin filaments thereby modulate the instability of these actin networks in a location-and dynamics-dependent manner. This regulatory role on the kinetics could be related to the formation of Arp2/3 puncta, which subsequently clustering other actin-binding proteins. Arp2/3 network was proposed to form in response to the mechanical cues (Papalazarou and Machesky, 2021; Skau and Waterman, 2015), and Arp2/3 generated actin network was also found to display different mechanical property (Bieling et al., 2016; Fritzsche et al., 2016; Muresan et al., 2022)(Jasnin et al., 2016). Since the basal membrane is connected with the substrate, it is perhaps logical that the branched actin network nucleated by Arp2/3 which is designed to produce pushing forces is less dominant here.

### Coordination between Arp2/3 and Formins

In the resting cortex of interphase HeLa cell, it was estimated about 10% of cortical actin were nucleated by formins (Fritzsche et al., 2016). The much higher relative contribution of formin mediated actin polymerization in actin wave in stimulated mast cell suggest the ratio of these nucleators can be readily tuned in a physiological context. Because of the challenges in visualizing their dynamics in unperturbed physiological context, the sequence of events leading to the interplay of these two types of nucleators in living cells remain unclear, and a number of different scenarios have been proposed. For instance, the leading-edge during migration involves the formation of Arp2/3 complex-dependent formation of lamellipodia, and formin nucleated filopodia composed of linear actin filaments. There, formin was proposed to act downstream of Arp2/3 by extending some barbed ends in the Arp2/3 nucleated dendritic tree and continue to elongate them (Svitkina et al., 2003). Consistent with this convergent elongation model, in a cell-free reconstitution system of filopodia formation, Diaphanous-related formin mDia2 was found to be recruited after Arp2/3 and actin (Lee et al., 2010). In vitro, formin mDia1 was recruited to Arp2/3 nucleated filaments and activated via a rocket launching mechanism (Breitsprecher et al., 2012; Cao et al., 2020). Conversely, formin has also been proposed to act upstream of Arp2/3, by providing the mother filaments where Arp2/3 can bind and generate new branches (Isogai et al., 2015; Yang et al., 2007).

Our characterization of actin wave nucleators confirms the crosstalk and synergy between these nucleators. The nucleators are recruited on the membrane and nucleate actin wave in a highly coordinated fashion, with a constant time-delay. Their mutual dependence and cooperation are also supported by the fact that without any external perturbation, these nucleators experiences spontaneous fluctuation of oscillation amplitudes in similar trend. In the Cryo-electron tomograph of actin waves in Dictyostelium, the presence of branched filaments and Arp2/3 at the junction were directly visualized (Jasnin et al., 2019). It was proposed that actin polymerization was initiated at the membrane by VASP and formins, followed by Arp2/3 complex accumulation and branch nucleation. This upstream role of formin relative to Arp2/3 is consistent with our observation that both FMNL1 and mDia3 are recruited 2-4 sec before Arp2/3. In addition, because CA-mutant of FMNL1 did not interfere with Arp2/3 recruitment, but a short CA-mutant lacking the actin binding FH1-FH2 domains (mini-FMNL1-V281E) negatively regulates Arp2/3 participation, it appears that Arp2/3 indeed benefited from FMNL1’s interaction with actin, directly or indirectly. On the other hand, Arp2/3 is highly clustered. FMNL1 and mDia3 are enriched in the waves but did not display highly contrasted clusters that colocalize with Arp2/3. The faint puncta of FMNL1 appear to be mutually exclusive compared to that of Arp2/3. In addition, consititutively active form of mDia3, with its FH domains and actin nucleation activity, inhibited Arp3 waves. In the Cryo-electron tomograph, the density of these junction and branched filaments were also relatively low compared to the density of total filaments. Thus, it seems likely that the majority of formin-nucleated filaments remain unbranched in its lifetime. Thus, the coordination and positive synergy most likely takes place at the level of actin networks with spatial proximity, but not always at the level of single filaments or branches.

### Competition between Arp2/3 and Formins

The regulation of Arp2/3 and formins also compete with each other at multiple time scale. When Arp2/3 is transiently inhibited, FMNL1 and mDia3’s surface recruitment increased. This antoganism occus within seconds. There is also a longer time scale competition because when the CA mutant of mDia3 or mini-FMNL1-V281E was expressed, Arp2/3’s recruitment was reduced or become non-detectable. Upregulation of formin activity upon Arp2/3 inhibition was previously observed in many contexts including in T cell (Murugesan et al., 2016). The most prominent model is that Arp2/3 and formin compete for a limiting pool of monomer (Burke et al., 2014; Henty-Ridilla and Goode, 2015; Rotty et al., 2015; Rotty and Bear, 2014; Suarez et al., 2015; Vitriol et al., 2015). Our data cannot be explained with this model because the monomer compitition model only explains the competition of formin and Arp2/3 activity (relative ratio of two types of filaments formed), but not the recruitment of nucleators themselves. In addition, the amplitude of formins or F-actin oscillation can easily fluctutate and we do not believe that actin waves are not operating in a regime where actin monomer is limiting. Instead, we favor a model where the limiting factor is short-lived and dynamically reset every cycle (20-30 sec). Our results are consistent with active Cdc42 being such limiting factor. Cdc42-dependent competition between Arp2/3 and formin operates through at least two mechanisms. First, the interactions between FMNL1 and Cdc42 are mutually exclusive with that of WASP and Cdc42. Second, Arp2/3 limits the amount of active Cdc42, which implies a negative feedback from Arp2/3 pathways. Arp2/3-dependent actin assembly could be particularlly important for endocytosis. Previously we have shown that ventral actin waves in mast cell are coupled with synchronized cycles of clathrin-mediated endocytosis (Yang et al., 2017). It is likely Arp2/3, together with endocytic fission events, limit the accumulation or dwell time of active Cdc42 on the plasma membrane, besides their well-established effector function. Because these inhibitory roles can only be revealed with experimental readout at fast time scale as well as moderate perturbation that does not completely inhibit actin turnover, they can be easily overlooked. The complexity and heterogeneity of actin wave therefore makes them an excellent model to dissect the interplay of actin regulators in the dynamic assembly of higher order structures. A comprehensive understanding of the intricate processes governing actin nucleation and assembly within these waves holds significant implications for unraveling how their organization and dynamics are matched with cell function in varied contexts (Bershadsky, 2004; Chhabra and Higgs, 2007; Skau and Waterman, 2015).

## Materials and Methods

### Cell Culture and transfection

Rat Basophilic Leukemia (RBL-2H3) cells (ATCC, CRL-2256) were maintained in monolayer cultures with MEM growth medium (Life Technologies, Carlsbad, CA) supplemented 20% heat inactivated Fetal Bovine Serum (Sigma-Aldrich, St Louis, MO) and harvested with TrypLE Express (Life Technologies, Carlsbad, CA). For transient transfections, 1.5 x 10^6^ cells were re-suspended in 10 μL of R buffer provided by the transfection kit with 1 μg of plasmid each, followed by two pulses at 1200 mV for 20 ms using Neon Transfection Electroporator (Life Technologies, Carlsbad, CA). After transfection, cells were plated at subconfluent densities in 35 mm glass-bottom Petri dishes (MatTek P35G-1.5-20-C). Transfected cells were maintained at a 37 °C humidified incubator overnight. The maximum number of cell passages for cells used in our experiments was limited to 30. For stimulation, transfected cells were sensitized overnight with anti-DNP IgE (Life Technologies, Carlsbad, CA) at 0.5 μg/mL. 80 ng/mL of DNP-BSA, a multivalent antigen that stimulates an antigen response, was added prior 1 h prior to imaging acquisition.

### Drug Treatment

For experiments using inhibitors, they were diluted from stock and added to the cells at the respective final concentrations: CK-666 (50-250 μM, Sigma-Aldrich, St Louis, MO), Cytochalasin-D (2 or 4 μM, Sigma-Aldrich, St Louis, MO), Wiskostatin (10 μM, Sigma-Aldrich, St Louis, MO). For experiments involving the chronic depletion of F-actin levels, transiently-transfected cells were incubated for 22 h in growth medium containing 2-4 μM cytochalasin-D before imaging. To allow recovery of the actin cytoskeleton, chronically-treated cells can be released into fresh growth medium for 2 h before imaging.

### Molecular Cloning and plasmids

The GFP-Arp3 (Wu Lab plasmid code A31), constructed by Matt Welch, was a gift from Dorothy Schafer (Schafer et al., 1998) (Human Arp3 was cloned into pEGFP-N1 using EcoRI and BamH1 sites). mCherry-Arp3 (Wu Lab plasmid code A36) was a gift from Michael Davidson (Addgene plasmid #54981). LifeAct-GFP (Wu Lab plasmid code A11) and LifeAct-mRuby (Wu Lab plasmid code A12) were gifts from Roland Wedlich-Solber (LifeAct was subcloned into pEGFP-N1 using EcoRI and BamHI sites. LifeAct-mRuby was generated by replacing GFP of LifeAct with mRuby (Fischer et al, 2006). For both LifeAct constructs, the linker GDPPVAT was generated between LifeAct and GFP or mRuby) (Riedl et al, 2008). FMNL1-GFP (Wu Lab plasmid code A98) was a gift from Michael Rosen (Seth et al, 2006) (FMNL1 was subcloned into a modified pCMV-Script vector containing C-terminal EGFP using BamHI and XhoI sites). GFP-CA-FMNL1 (L1062D mutation) (Wu Lab plasmid code A98F), GFP-FMNL1CT (aa 612-1094) (Wu Lab plasmid code A98C), GFP-mini-FMNL1 (Wu Lab plasmid code A98E), GFP-mini-FMNL1-V281E (Wu Lab plasmid code A98G), GFP-mini-FMNL1-T126D (Wu Lab plasmid code A98H) were gifts from Michael Rosen (Seth et al, 2006). FMNL1-iFP (Wu Lab plasmid code A98K) was generated in our lab and subcloned using XhoI and BamHI sites from FMNL1-GFP where GFP was replaced with iFP fluorescent tag. GFP-mDia3 (Wu Lab plasmid code A99) was constructed by Shuh Narumiya through subcloning the fragment encoding mDia3 from pCR vector into pEGFP vector using AccI and BamHI sites (Yasuda et al, 2004). mCherry-mDia3 (Wu Lab plasmid code A99C) was generated in our lab and subcloned using SacI and BamHI sites from GFP-mDia3 into mCherry-C1 backbone vector. mCherry-CA-mDia3 (aa 1-1053) (Wu Lab plasmid code A99B) was generated by overlap extension PCR for the deletion of the DAD domain from the full length-mDia3 encoded by mCherry-mDia3. PM-mScarlet (Wu Lab plasmid code M02B) was generating from subcloning of the mScarlet fluorescent tag using PCR from PMScarlet-H C1 (Bindels et al, 2016) into PM-miRFP670 where miRFP670 was replaced with the mScarlet fluorescent tag. Cdc42 BD-GFP (Wu Lab plasmid code G011), a biosensor for detecting endogenous active Cdc42 generated by cloning bacterial expressing Cdc42 activity sensor pET23-CBD-(-PP)-EGFP (Nalbant et al, 2004) into pEGFP-N1 vector with restriction sites XhoI and BamHI. For experiments using Cdc42 BD-GFP, a stable cell line was generated. Cdc42 BD-GFP plasmid was transfected into RBL-2H3 cells for 24 h, selected in MEM containing 20% FBS and 0.5 mg/ml selection antibiotic G418 and FACS sorted.

### Microscopy

Before imaging, medium in the dishes were replaced with fresh normal growth medium (except for experiments involving cytochalasin-D) and transferred to a heated microscope stage (Live Cell Instrument, Seoul, South Korea) maintained at 37 °C throughout the experiments. For TIRFM imaging of live cells’ cortical dynamics, a Nikon Ti-E inverted microscope (Nikon, Shinagawa, Tokyo) was used. The microscope was equipped with a perfect focus system that prevents focus drift, an iLAS2 motorized TIRF illuminator (Roper Scientific, Evry Cedex, France) and either an Evolve 512 EMCCD camera (Photometrics, Tucson, AZ) (16 bit, pixel size 16 μm) or Prime95b sCMOS camera (Photometrics, Tucson, AZ) (16 bit, pixel size 11 μm). All images were acquired using objective lenses from Nikon’s CFI Apochromat TIRF Series (100xH N.A. 1.49 Oil; 60xH N.A. 1.49 Oil). Multi-channel imaging of samples was achieved by the sequential excitation with 491 nm (100 mW), 561 nm (100 mW) and 642 nm (100 mW) lasers, reflected from a quad-bandpass dichroic mirror (Di01-R405/488/561/635, Semrock, Rochester, NY) located on a Ludl emission filter wheel (Carl Zeiss AG, Oberkochen, Germany). The microscope was controlled by a MetaMorph software (Version 7.8.6.0) (Molecular Devices, LLC, Suunyvale, CA). For sequential TIRF/Differential interference contrast (DIC) imaging, channel transition was controlled by Multiple Dimension Acquisition module in Metamorph and a Normaski primary prism was inserted in condenser. During image acquisition, the samples were maintained at 37 °C with an on-stage incubator system (Live Cell Instrument, Seoul, South Korea). To ensure cell viability during long term imaging lasting more than one hour, spent media was replaced with fresh media. In addition, 5% humidified CO_2_ was supplied. All image sequences were acquired at time intervals of 1-3 sec, except for extended acquisition durations lasting over 45 min, which were acquired at time intervals of 4 sec.

### Image analysis

Post-acquisition image analyses were performed by either by Fiji (Schindelin et al, 2012) or MATLAB (The MathWorks, Inc., Natick, MA). Kymographs were generated using the ‘Reslice’ tool. Depending on intensity of the fluorescent markers, ‘average’ projection filters (average of 10 frames) and background subtractions may be applied to enhance presentation. For consistency, the same image processing was applied across all channels for kymographs generated from multi-color imaging and applied identically across different conditions that were directly compared to each other. Sequential montages depicting the wave front were generated using the ‘Make Montage’ tool. For intensity profiles, a region of interest with a 20 x 20 pixels size was used for all movies. The intensity data points generated by Fiji were normalized and plotted by Matlab. To illustrate phase differences between proteins in wave propagation, multiple cycles of their intensity fluctuations were aligned. The solid lines represent the mean intensities and shaded region represent the standard deviations of the intensities. To obtain the relative intensity, the intensity values were normalized by the background substracted average intensity within the individual cell. Cells positive of waves was determined by visual impression and confirmed using the built-in Fast Fourier Transform function in Matlab. Cells that gave an output of a single FFT peak were deemed positive of waves and cells without the FFT peak were deemed negative of waves. Wavelet analysis, auto-correlation, cross-correlation, phase plots were performed using MATLAB. Custom codes used for analysis are deposited on Github (https://github.com/min-wu-lab/mmo-analysis, https://doi.org/10.5281/zenodo.8083400) and is publicly available.

### Statistical analysis

All statistical analysis were performed by Prism 7 (GraphPad Software, Inc, La Jolla, CA). For comparison between control and perturbed samples, unpaired two-tailed student’s t-test was performed. For multiple comparisons of frequencies in Figure 3D, a one-way ANOVA was performed. Unless otherwise stated, error bars for all data shown represents mean ± S.E.M.

## Resource availability

### Lead contact

Further information and requests for resources and reagents should be directed to and will be fulfiled by the lead contact, Min Wu.

### Materials availability

All unique/stable reagents generated in this study are available from the lead contact without restriction.

### Data and code availability

All data reported in this paper will be shared by the lead contact upon request. The code used to generate the figures has been previously documented and is openly accessible on GitHub, publicity available since the publication date (https://doi.org/10.5281/zenodo.8083400). Source data used for analysis in this paper is uploaded to Mendeley data (DOI: 10.17632/kt34mc26k3.1). Any additional information required to re-analyze the data reported in this paper is available from the lead contact upon request.

## Supporting information

Video 1, Arp2/3 complex assembles in actin waves

Video 2, Formin FMNL1 assembles before Arp2/3 complex

Video 3, coordinated waves of the constitutively-active FMNL1 and Arp2/3 complex

Video 4, punctate waves of FMNL1CT co-localizes with Arp2/3 puncta

Video 5, Cdc42 waves de-coupled from actin waves

Video 6, Cdc42 oscillations with or without actin waves

Video 7, CA-FMNL1 oscillations de-coupled from actin waves

Video 8, Arp2/3 inhibition enhanced FMNL1 waves

Video 9, Arp2/3 inhibition enhanced mDia3 waves

Video 10, Arp2/3 inhibition enhanced Cdc42 waves

## Acknowledgements

We thank Michael Rosen, Dorothy Schafer, Roland Wedlich-Solber, Shuh Narumiya and Alexander Bershadsky for providing plasmids used in our study; Jeffery Yong and Guo Su for technical assistance; former and current members of the Wu lab and Pekka Lappalainen for discussions. X.L.C was affiliated with the Department of Cell Biology, Yale University School of Medicine, USA when the experiments were performed and is currently affiliated with the HiLIFE Insititute of Biotechnology, University of Helsinki, Finland. This work is supported by Yale University startup grant (M.W.) and National Institute Of General Medical Sciences of the National Institutes of Health under Award Number R01GM151344 (M.W.). The content is solely the responsibility of the authors and does not necessarily represent the official views of the National Institutes of Health.

## Author contributions

M.W. and M.S. conceptualized the project. X.L.C. performed most of the experiments and analyzed the results with contributions from C.S.T., and X.J.X. X.L.C., C.S.T., and M.W., designed the experiments and interpreted the results. S.X. discovered the cytochalasin-D treatment and X.L.C. optimized the protocol. X.L.C., and M.W., wrote the manuscript with contributions from C.S.T., and X.J.X. M.W. designed the MATLAB codes for data analysis. X.J.X developed the python analysis software. X.W. contributed to the structural models. M.W. supervised the project.

---

**Figure S1.**
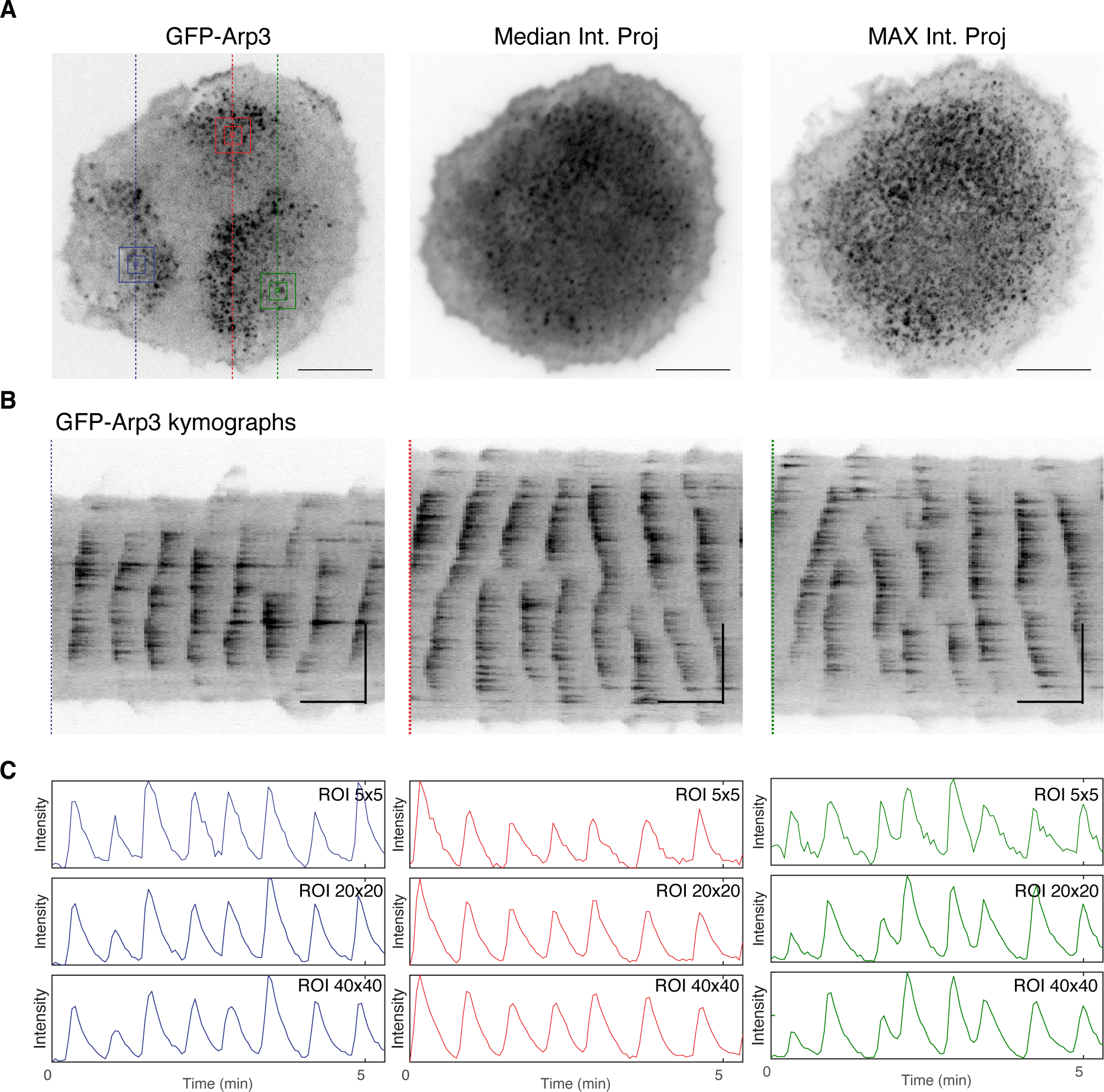
Qualitative and quantitative post-acquisition analysis of cell with arbitrarily chosen regions. **(A)** Representative micrographs of a cell expressing GFP-Arp3. Median intensity- and maximum intensity projections of timelapse images of the representative cell exhibiting waves of GFP-Arp3 puncta. **(B)** Kymographs re-sliced from 10 continuous frames with average projection from three arbitrarily chosen region of the cell (left: blue; middle: red; right: green) to demonstrate cycles of wave propagation over time. **(C)** Fluorescence intensity plots derived from three arbitrarily chosen regions of ROIs of different pixel sizes (5×5 pixels; 20×20 pixels; 40×40 pixels).

**Figure S2.**
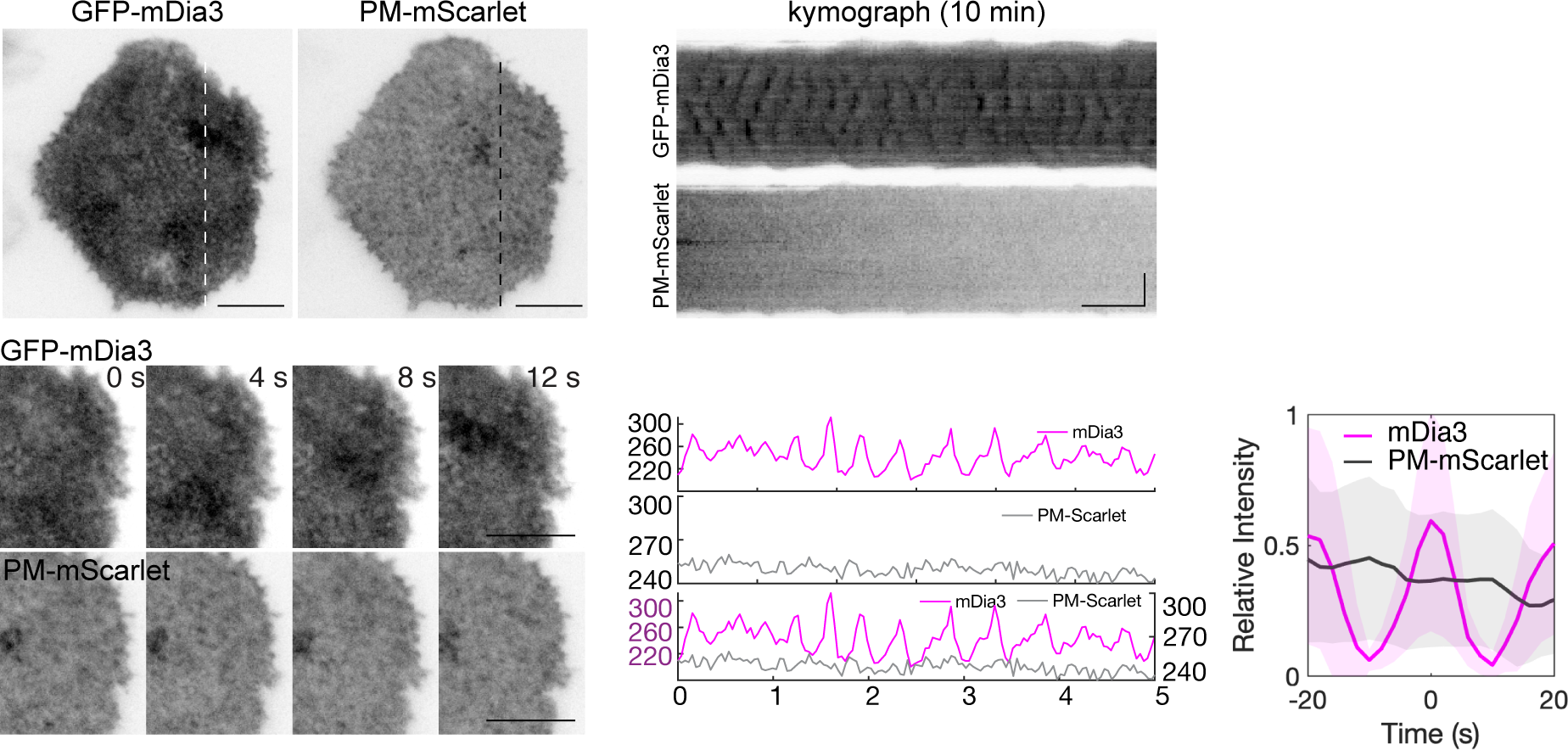
Cortical waves of GFP-mDia3 compared with a plasma membrane marker. Representative micrographs and kymographs of a cell co-expressing GFP-mDia3 and PM-mScarlet. Fluorescence intensity plots of a cell co-expressing GFP mDia3 (magenta) and PM-mScarlet (grey). Fluorescence intensity profile of PM-mScarlet (grey) aligned with respect to GFP-mDia3 (magenta). Horizontal scale bars in micrographs: 10 µm. Vertical scale bars in kymographs: 10 µm. Horizontal scale bars in kymographs: 1 min.

## Video Legends

**Video 1. Arp2/3 complex assembles in actin waves**

Coordinated waves of Arp2/3 and F-actin puncta assemble and disassemble at the cortex of a RBL-2H3 cell. TIRFM video of a cell expressing GFP-Arp3 (green) and LifeAct-mRuby (magenta) was acquired at 3 sec per frame. Scale bar 10 µm.

**Video 2. Formin assembles before Arp2/3 complex**

Diffuse waves of FMNL1 trailed by waves of Arp2/3 puncta at the cortex of a RBL-2H3 cell. TIRFM video of a cell expressing FMNL1-GFP (green) and mCherry-Arp3 (magenta) was acquired at 4 sec per frame. Scale bar 10 µm.

**Video 3. Constitutively-active mutant of FMNL1 and Arp2/3 assembles in wave**

Waves of the constitutively-active FMNL1 precedes Arp2/3 complex at the cortex of a RBL-2H3 cell. TIRFM video of a cell expressing GFP-FMNL1-L1062D (green) and mCherry-Arp3 (magenta) was acquired at 3 sec per frame. Playback rate: 6 frames per sec, 18× real time. Scale bar 10 µm.

**Video 4. Punctate waves of FMNL1 co-localizes with Arp2/3 puncta**

C-termini fragment of FMNL1 exhibits clusters of puncta that co-localizes with Arp2/3 puncta at the cortex of a RBL-2H3 cell. TIRFM video of a cell expressing GFP-FMNL1CT (green) and mCherry-Arp3 (magenta) was acquired at 3 sec per frame. Playback rate: 6 frames per sec, 18× real time. Scale bar 10 µm.

**Video 5. Waves of Cdc42 de-coupled from F-actin waves**

Waves of active Cdc42 propagates when F-actin waves are abolished by the chronic treatment of cytochalasin. TIRFM video of a cell expressing Cdc42 BD-GFP and LifeAct-mRuby was acquired at 3 sec per frame. Playback rate: 20 frames per sec, 60× real time. Scale bar 10 µm.

**Video 6. Cdc42 oscillations with or without F-actin waves**

Oscillatory periodicity of active Cdc42 waves slows down when F-actin waves are abolished by the chronic treatment of cytochalasin. TIRFM video of an untreated cell (top) expressing Cdc42 BD-GFP and LifeAct-mRuby was acquired at 3 sec per frame. Playback rate: 8 frames per sec. TIRFM video of a cell chronically treated by cytochalasin (bottom) expressing Cdc42 BD-GFP and LifeAct-mRuby was acquired at 4 sec per frame. Playback rate: 6 frames per sec. 24× real time. Scale bar 10 µm.

**Video 7. Waves of constitutively-active FMNL1 de-coupled from F-actin waves**

Waves exhibited by the constitutively-active FMNL1 propagates when F-actin waves are abolished by the chronic treatment of cytochalasin. TIRFM video of a cell expressing GFP-FMNL1-L1062D and LifeAct-mRuby was acquired at 3 sec per frame. Playback rate: 20 frames per sec, 60× real time. Scale bar 10 µm.

**Video 8. FMNL1 waves enhanced after Arp2/3 inhibition**

Fluorescence intensities for the cortical waves of FMNL1 increased while most of the Arp2/3 puncta disappeared when cell was treated by CK666 at 50-150µM. TIRFM video of a cell expressing FMNL1-GFP and mCherry-Arp3 was acquired at 3 sec per frame. Playback rate: 20 frames per sec, 60× real time. Scale bar 10 µm.

**Video 9. mDia3 waves enhanced after Arp2/3 inhibition**

Fluorescence intensities for the cortical waves of mDia3 increased while most of the Arp2/3 puncta disappeared when cell was treated by CK666 at 50-150µM. TIRFM video of cells expressing GFP-mDia3 and mCherry-Arp3 was acquired at 3 sec per frame. Playback rate: 20 frames per sec, 60× real time. Scale bar 10 µm.

**Video 10. Arp2/3 inhibition enhanced Cdc42 waves**

Fluorescence intensities for the cortical waves of active Cdc42 increased while most of the Arp2/3 puncta disappeared when cell was treated by CK666 at 50 µM. TIRFM video of cells expressing Cdc42 BD-GFP and mCherry-Arp3 was acquired at 3 sec per frame. Playback rate: 20 frames per sec, 60× real time. Scale bar 10 µm.

## Notes

https://doi.org/10.5281/zenodo.8083400

DOI: 10.17632/kt34mc26k3.1

### Competing Interest Statement

The authors have declared no competing interest.

## Reference

Abdul-Manan N, Aghazadeh B, Liu GA, Majumdar A, Ouerfelli O, Siminovitch KA, Rosen MK. 1999. Structure of Cdc42 in complex with the GTPase-binding domain of the “Wiskott–Aldrich syndrome” protein. Nature 399:379–383.

Bement WM, Leda M, Moe AM, Kita AM, Larson ME, Golding AE, Pfeuti C, Su KC, Miller AL, Goryachev AB, Von Dassow G. 2015. Activator-inhibitor coupling between Rho signalling and actin assembly makes the cell cortex an excitable medium. Nat Cell Biol 17:1471–1483.

Bershadsky A. 2004. Magic touch: how does cell-cell adhesion trigger actin assembly? Trends Cell Biol 14:1–5.

Bieling P, Li T De, Weichsel J, McGorty R, Jreij P, Huang B, Fletcher DA, Mullins RD. 2016. Force Feedback Controls Motor Activity and Mechanical Properties of Self-Assembling Branched Actin Networks. Cell 164:115–127.

Bolado-Carrancio A, Rukhlenko OS, Nikonova E, Tsyganov MA, Wheeler A, Garcia-Munoz A, Kolch W, Kriegsheim A von, Kholodenko BN. 2020. Periodic propagating waves coordinate rhogtpase network dynamics at the leading and trailing edges during cell migration. Elife 9:1–34.

Breitsprecher D, Jaiswal R, Bombardier JP, Gould CJ, Gelles J, Goode BL. 2012. Rocket Launcher Mechanism of Collaborative Actin Assembly Defined by Single-Molecule Imaging. Science (1979) 336:1164–1167.

Bretschneider T, Anderson K, Ecke M, Müller-Taubenberger A, Schroth-Diez B, Ishikawa-Ankerhold HC, Gerisch G. 2009. The three-dimensional dynamics of actin waves, a model of cytoskeletal self-organization. Biophys J 96:2888–2900.

Bretschneider T, Diez S, Anderson K, Heuser J, Clarke M, Müller-Taubenberger A, Köhler J, Gerisch G. 2004. Dynamic Actin Patterns and Arp2/3 Assembly at the Substrate-Attached Surface of Motile Cells. Current Biology 14:1–10.

Burke TA, Christensen JR, Barone E, Suarez C, Sirotkin V, Kovar DR. 2014. Homeostatic actin cytoskeleton networks are regulated by assembly factor competition for monomers. Current Biology 24:579–585.

Campellone KG, Welch MD. 2010. A nucleator arms race: Cellular control of actin assembly. Nat Rev Mol Cell Biol 11:237–251.

Cao L, Yonis A, Vaghela M, Barriga EH, Chugh P, Smith MB, Maufront J, Lavoie G, Méant A, Ferber E, Bovellan M, Alberts A, Bertin A, Mayor R, Paluch EK, Roux PP, Jégou A, Romet-Lemonne G, Charras G. 2020. SPIN90 associates with mDia1 and the Arp2/3 complex to regulate cortical actin organization. Nat Cell Biol 22:803–814.

Chhabra ES, Higgs HN. 2007. The many faces of actin: matching assembly factors with cellular structures. Nat Cell Biol 9:1110–1121.

Dehapiot B, Clément R, Alégot H, Gazsó-Gerhát G, Philippe JM, Lecuit T. 2020. Assembly of a persistent apical actin network by the formin Frl/Fmnl tunes epithelial cell deformability. Nat Cell Biol 22:791– 802.

Dimitracopoulos A, Srivastava P, Chaigne A, Win Z, Shlomovitz R, Lancaster OM, Le Berre M, Piel M, Franze K, Salbreux G, Baum B. 2020. Mechanochemical Crosstalk Produces Cell-Intrinsic Patterning of the Cortex to Orient the Mitotic Spindle. Current Biology 30:3687–3696.

Ecke M, Prassler J, Tanribil P, Müller-Taubenberger A, Körber S, Faix J, Gerisch G. 2020. Formins specify membrane patterns generated by propagating actin waves. Mol Biol Cell 31:373–385.

Flemming S, Font F, Alonso S, Beta C. 2020. How cortical waves drive fission of motile cells. Proc Natl Acad Sci USA 117:6330–6338.

Fritzsche M, Erlenkämper C, Moeendarbary E, Charras G, Kruse K. 2016. Actin kinetics shapes cortical network structure and mechanics. Sci Adv 2:1–12.

Fukushima S, Matsuoka S, Ueda M. 2019. Excitable dynamics of Ras triggers spontaneous symmetry breaking of PIP3 signaling in motile cells. J Cell Sci 132:1–12.

Gerisch G. 2010. Self-organizing actin waves that simulate phagocytic cup structures. PMC Biophys 3:1–9.

Gerisch G, Ecke M, Schroth-Diez B, Gerwig S, Engel U, Maddera L, Clarke M. 2009. Self-organizing actin waves as planar phagocytic cup structures. Cell Adh Migr 3:373–382.

Gerisch G, Ecke M, Wischnewski D, Schroth-Diez B. 2011. Different modes of state transitions determine pattern in the Phosphatidylinositide-Actin system. BMC Cell Biol 12:1–15.

Graessl M, Koch J, Calderon A, Kamps D, Banerjee S, Mazel T, Schulze N, Jungkurth JK, Patwardhan R, Solouk D, Hampe N, Hoffmann B, Dehmelt L, Nalbant P. 2017. An excitable Rho GTPase signaling network generates dynamic subcellular contraction patterns. Journal of Cell Biology 216:4271–4285.

Henty-Ridilla JL, Goode BL. 2015. Global resource distribution: Allocation of actin building blocks by profilin. Dev Cell 32:5–6.

Hetrick B, Han MS, Helgeson LA, Nolen BJ. 2013. Small molecules CK-666 and CK-869 inhibit actin-related protein 2/3 complex by blocking an activating conformational change. Chem Biol 20:701–712.

Honda G, Saito N, Fujimori T, Hashimura H, Nakamura MJ, Nakajima A, Sawai S. 2021. Microtopographical guidance of macropinocytic signaling patches. Proc Natl Acad Sci USA 118:1–11.

Hörning M, Bullmann T, Shibata T. 2021. Local Membrane Curvature Pins and Guides Excitable Membrane Waves in Chemotactic and Macropinocytic Cells - Biomedical Insights From an Innovative Simple Model. Front Cell Dev Biol 9:1–19.

Huang CH, Tang M, Shi C, Iglesias PA, Devreotes PN. 2013. An excitable signal integrator couples to an idling cytoskeletal oscillator to drive cell migration. Nat Cell Biol 15:1307–1316.

Huang YA, Hsu CH, Chiu HC, Hsi PY, Ho CT, Lo WL, Hwang E. 2020. Actin waves transport RanGTP to the neurite tip to regulate non-centrosomal microtubules in neurons. J Cell Sci 133:1–13.

Isogai T, van der Kammen R, Leyton-Puig D, Kedziora KM, Jalink K, Innocenti M. 2015. Initiation of lamellipodia and ruffles involves cooperation between mDia1 and the Arp2/3 complex. J Cell Sci 128:3796–3810.

Jasnin M, Beck F, Ecke M, Fukuda Y, Martinez-Sanchez A, Baumeister W, Gerisch G. 2019. The Architecture of Traveling Actin Waves Revealed by Cryo-Electron Tomography. Structure 27:1211–1223.

Kim AS, Kakalis LT, Abdul-Manan N, Liu GA, Rosen MK. 2000. Autoinhibition and activation mechanisms of the Wiskott±Aldrich syndrome protein. Nature 404:151–158.

Kühn S, Erdmann C, Kage F, Block J, Schwenkmezger L, Steffen A, Rottner K, Geyer M. 2015. The structure of FMNL2-Cdc42 yields insights into the mechanism of lamellipodia and filopodia formation. Nat Commun 6:1–14.

Lam Hui K, Kwak SI, Upadhyaya A. 2014. Adhesion-dependent modulation of actin dynamics in Jurkat T cells. Cytoskeleton 71:119–135.

Lappalainen P, Kotila T, Jégou A, Romet-Lemonne G. 2022. Biochemical and mechanical regulation of actin dynamics. Nat Rev Mol Cell Biol 23:836–852.

Lee K, Gallop JL, Rambani K, Kirschner MW. 2010. Self-Assembly of Filopodia-Like Structures on Supported Lipid Bilayers. Science (1979) 329:1341–1345.

Li X, Miao Y, Pal DS, Devreotes PN. 2020. Excitable networks controlling cell migration during development and disease. Semin Cell Dev Biol 100:133–142.

Lutton JE, Coker HLE, Paschke P, Munn CJ, King JS, Bretschneider T, Kay RR. 2023. Formation and closure of macropinocytic cups in Dictyostelium. Current Biology 33:3083–3096.

Machesky LM, Dyche Mullins R, Higgs HN, Kaiser DA, Blanchoin L, May RC, Hall ME, Pollard TD. 1999. Scar, a WASp-related protein, activates nucleation of actin filaments by the Arp23 complex. Proc Natl Acad Sci U S A 96:3739–3744.

Machesky LM, Insall RH. 1998. Scar1 and the related Wiskott-Aldrich syndrome protein, WASP, regulate the actin cytoskeleton through the Arp2/3 complex. Current Biology 8:1347–1356.

Maître JL, Niwayama R, Turlier H, Nedelec F, Hiiragi T. 2015. Pulsatile cell-autonomous contractility drives compaction in the mouse embryo. Nat Cell Biol 17:849–855.

Masters TA, Sheetz MP, Gauthier NC. 2016. F-actin waves, actin cortex disassembly and focal exocytosis driven by actin-phosphoinositide positive feedback. Cytoskeleton 73:180–196.

Miao Y, Bhattacharya S, Banerjee T, Abubaker-Sharif B, Long Y, Inoue T, Iglesias PA, Devreotes PN. 2019. Wave patterns organize cellular protrusions and control cortical dynamics. Mol Syst Biol 15:1–20.

Michaud A, Leda M, Swider ZT, Kim S, He J, Landino J, Valley JR, Huisken J, Goryachev AB, Von Dassow G, Bement WM. 2022. A versatile cortical pattern-forming circuit based on Rho, F-actin, Ect2, and RGA-3/4. Journal of Cell Biology 221:1–21.

Michaux JB, Robin FB, McFadden WM, Munro EM. 2018. Excitable RhoA dynamics drive pulsed contractions in the early C. Elegans embryo. Journal of Cell Biology 217:4230–4252.

Miki H, Sasaki T, Takai Y, Takenawa T. 1998. Induction of filopodium formation by a WASP-related actin-depolymerizing protein N-WASP. Nature 391:93–96.

Mitsushima M, Aoki K, Ebisuya M, Matsumura S, Yamamoto T, Matsuda M, Toyoshima F, Nishida E. 2010. Revolving movement of a dynamic cluster of actin filaments during mitosis. Journal of Cell Biology 191:453–462.

Moore AS, Coscia SM, Simpson CL, Ortega FE, Wait EC, Heddleston JM, Nirschl JJ, Obara CJ, Guedes-Dias P, Boecker CA, Chew TL, Theriot JA, Lippincott-Schwartz J, Holzbaur ELF. 2021. Actin cables and comet tails organize mitochondrial networks in mitosis. Nature 591:659–664.

Moore AS, Wong YC, Simpson CL, Holzbaur ELF. 2016. Dynamic actin cycling through mitochondrial subpopulations locally regulates the fission-fusion balance within mitochondrial networks. Nat Commun 7:1–13.

Mullins RD, Pollard TD. 1999. Rho-family GTPases require the Arp2/3 complex to stimulate actin polymerization in Acanthamoeba extracts. Current Biology 9:405–415.

Muresan CG, Sun ZG, Yadav V, Tabatabai AP, Lanier L, Kim JH, Kim T, Murrell MP. 2022. F-actin architecture determines constraints on myosin thick filament motion. Nat Commun 13:1–16.

Murugesan S, Hong J, Yi J, Li D, Beach JR, Shao L, Meinhardt J, Madison G, Wu X, Betzig E, Hammer JA. 2016. Formin-generated actomyosin arcs propel t cell receptor microcluster movement at the immune synapse. Journal of Cell Biology 215:383–399.

Nishikawa M, Naganathan SR, Jü Licher F, Grill SW. 2017. Controlling contractile instabilities in the actomyosin cortex. Elife 6:1–21.

Özgüç Ö, de Plater L, Kapoor V, Tortorelli AF, Clark AG, Maître JL. 2022. Cortical softening elicits zygotic contractility during mouse preimplantation development. PLoS Biol 20:1–23.

Papalazarou V, Machesky LM. 2021. The cell pushes back: The Arp2/3 complex is a key orchestrator of cellular responses to environmental forces. Curr Opin Cell Biol 68:37–44.

Rohatgi R, Ho H-YH, Kirschner MW. 2000. Mechanism of N-WASP Activation by CDC42 and Phosphatidylinositol 4,5-bisphosphate. J Cell Biol 150:1299–1309.

Rohatgi R, Ma L, Hiroaki M, Lopez M, Kirchhausen T, Takenawa T, Kirschner MW. 1999. The Interaction between N-WASP and the Arp2/3 Complex Links Cdc42-Dependent Signals to Actin Assembly. Cell 97:221–231.

Rotty JD, Bear JE. 2014. Competition and collaboration between different actin assembly pathways allows for homeostatic control of the actin cytoskeleton. Bioarchitecture 5:27–34.

Rotty JD, Brighton HE, Craig SL, Asokan SB, Cheng N, Ting JP, Bear JE. 2017. Arp2/3 Complex Is Required for Macrophage Integrin Functions but Is Dispensable for FcR Phagocytosis and In Vivo Motility. Dev Cell 42:498–513.

Rotty JD, Wu C, Haynes EM, Suarez C, Winkelman JD, Johnson HE, Haugh JM, Kovar DR, Bear JE. 2015. Profilin-1 Serves as a gatekeeper for actin assembly by Arp2/3-Dependent and - Independent pathways. Dev Cell 32:54–67.

Saito N, Sawai S. 2021. Three-dimensional morphodynamic simulations of macropinocytic cups. iScience 24:1–22.

Schroth-Diez B, Gerwig S, Ecke M, Heger R, Diez S, Gerisch G. 2009. Propagating waves separate two states of actin organization in living cells. HFSP J 3:412–427.

Seth A, Otomo C, Rosen MK. 2006. Autoinhibition regulates cellular localization and actin assembly activity of the diaphanous-related formins FRLα and mDia1. Journal of Cell Biology 174:701–713.

Skau CT, Waterman CM. 2015. Specification of Architecture and Function of Actin Structures by Actin Nucleation Factors. Annu Rev Biophys 44:285–310.

Stankevicins L, Ecker N, Terriac E, Maiuri P, Schoppmeyer R, Vargas P, Lennon-Duménil A-M, Piel M, Qu B, Hoth M, Kruse K, Lautenschläger F. 2020. Deterministic actin waves as generators of cell polarization cues. Proc Natl Acad Sci USA 117:826–835.

Su M, Zhuang Y, Miao X, Zeng Y, Gao W, Zhao W, Wu M. 2020. Comparative Study of Curvature Sensing JC 20230912jerrymwMediated by F-BAR and an Intrinsically Disordered Region of FBP17. iScience 23:1–16.

Suarez C, Carroll RT, Burke TA, Christensen JR, Bestul AJ, Sees JA, James ML, Sirotkin V, Kovar DR. 2015. Profilin regulates F-Actin network homeostasis by favoring formin over Arp2/3 complex. Dev Cell 32:43–53.

Suraneni P, Rubinstein B, Unruh JR, Durnin M, Hanein D, Li R. 2012. The Arp2/3 complex is required for lamellipodia extension and directional fibroblast cell migration. Journal of Cell Biology 197:239–251.

Svitkina TM, Bulanova EA, Chaga OY, Vignjevic DM, Kojima S ichiro, Vasiliev JM, Borisy GG. 2003. Mechanism of filopodia initiation by reorganization of a dendritic network. Journal of Cell Biology 160:409–421.

Swider ZT, Michaud A, Leda M, Landino J, Goryachev AB, Bement WM. 2022. Cell cycle and developmental control of cortical excitability in Xenopus laevis. Mol Biol Cell 33:1–13.

Tan TH, Liu J, Miller PW, Tekant M, Dunkel J, Fakhri N. 2020. Topological turbulence in the membrane of a living cell. Nat Phys 16:657–662.

Tong CS, Su M, Sun H, Chua X Le, Guo S, Ramaraj RS, Ong NPW, Lee AG, Miao Y, Wu M. 2023a. Collective dynamics of formin and microtubule and its crosstalk mediated by FHDC1. bioRxiv 1–28.

Tong CS, Xŭ XJ, Wu M. 2023b. Periodicity, mixed-mode oscillations, and multiple timescales in a phosphoinositide-Rho GTPase network. Cell Rep 42:1–18.

Vicker MG. 2002. Eukaryotic cell locomotion depends on the propagation of self-organized reaction-diffusion waves and oscillations of actin filament assembly. Exp Cell Res 275:54–66.

Vitriol EA, McMillen LM, Kapustina M, Gomez SM, Vavylonis D, Zheng JQ. 2015. Two functionally distinct sources of actin monomers supply the leading edge of lamellipodia. Cell Rep 11:433–445.

Weiner OD, Marganski WA, Wu LF, Altschuler SJ, Kirschner MW. 2007. An actin-based wave generator organizes cell motility. PLoS Biol 5:2053–2063.

Wigbers MC, Tan TH, Brauns F, Liu J, Swartz SZ, Frey E, Fakhri N. 2021. A hierarchy of protein patterns robustly decodes cell shape information. Nat Phys 17:578–584.

Wollman R, Meyer T. 2012. Coordinated oscillations in cortical actin and Ca2+ correlate with cycles of vesicle secretion. Nat Cell Biol 14:1261–1269.

Wu C, Asokan SB, Berginski ME, Haynes EM, Sharpless NE, Griffith JD, Gomez SM, Bear JE. 2012. Arp2/3 Is Critical for Lamellipodia and Response to Extracellular Matrix Cues but Is Dispensable for Chemotaxis. Cell 148:973–987.

Wu M, Wu X, De Camilli P. 2013. Calcium oscillations-coupled conversion of actin travelling waves to standing oscillations. Proc Natl Acad Sci U S A 110:1339–1344.

Wu Z, Su M, Tong C, Wu M, Liu J. 2018. Membrane shape-mediated wave propagation of cortical protein dynamics. Nat Commun 9:1–12.

Xiao S, Tong C, Yang Y, Wu M. 2017. Mitotic Cortical Waves Predict Future Division Sites by Encoding Positional and Size Information. Dev Cell 43:493–506.

Xiong D, Xiao S, Guo S, Lin Q, Nakatsu F, Wu M. 2016. Frequency and amplitude control of cortical oscillations by phosphoinositide waves. Nat Chem Biol 12:159–166.

Yang C, Czech L, Gerboth S, Kojima SI, Scita G, Svitkina T. 2007. Novel roles of formin mDia2 in lamellipodia and filopodia formation in motile cells. PLoS Biol 5:2624–2645.

Yang Y, Xiong D, Pipathsouk A, Weiner OD, Wu M. 2017. Clathrin Assembly Defines the Onset and Geometry of Cortical Patterning. Dev Cell 43:507–521.

Yao B, Donoughe S, Michaux J, Munro E. 2022. Modulating RhoA effectors induces transitions to oscillatory and more wavelike RhoA dynamics in Caenorhabditis elegans zygotes. Mol Biol Cell 33:1–16.

Zhan H, Bhattacharya S, Cai H, Iglesias PA, Huang CH, Devreotes PN. 2020. An Excitable Ras/PI3K/ERK Signaling Network Controls Migration and Oncogenic Transformation in Epithelial Cells. Dev Cell 54:608–623.

